# Changes in Urinary Proteome Post-Translational Modifications Following Short-Term Intake of Specific Magnesium, Calcium, Zinc, and Iron Supplements in Rats

**DOI:** 10.1101/2025.09.17.676746

**Authors:** Yan Su, Ziyun Shen, Youhe Gao

## Abstract

Magnesium, calcium, zinc, and iron are essential mineral elements indispensable for maintaining physiological functions in mammals. This study investigated changes in post-translational modifications (PTMs) of the urinary proteome in Sprague-Dawley (SD) rats of different ages following short-term intragastric administration of magnesium L-threonate (MgT), calcium gluconate, zinc gluconate, and polysaccharide-iron complex. The results revealed PTM alterations in the urinary proteins of rats in all mineral supplement groups, with the most pronounced changes observed in the polysaccharide-iron complex group. Notably, rats of different ages exhibited distinct responses to the same mineral supplement. The proteins carrying differential PTMs were involved in a wide range of biological functions, and several of them, including serotransferrin and beta-2-microglobulin, are closely associated with mineral metabolism. This study explores the functional changes in urinary proteins from a PTM perspective following short-term mineral supplementation in rats, providing a novel viewpoint for investigating the physiological functions of mineral elements.

## 1 Introduction

Magnesium, calcium, zinc, and iron are essential mineral elements indispensable for maintaining physiological functions in mammals. Magnesium ions serve as critical cofactors for over 300 enzymes, participating in energy metabolism, DNA repair, and cellular signal transduction. By regulating ATPase activity, magnesium notably influences cellular energy supply; its deficiency can lead to metabolic disorders and neurological dysfunction^[1]^. Calcium is not only a major component of bones and teeth but also acts as a key second messenger in neural transmission, muscle contraction, and hormone secretion^[2]^. Zinc functions as a transcription factor activator and a component of antioxidant enzymes. It helps maintain cellular homeostasis by regulating immune responses and promoting protein synthesis; zinc deficiency may result in growth retardation and impaired immune function^[3]^. Iron, as a core component of hemoglobin and cytochromes, is essential for oxygen transport and redox reactions. Its metabolic imbalance is closely associated with anemia and oxidative stress^[4]^. In recent years, the potential of mineral supplements in preventing and treating metabolic diseases has gained increasing attention. Previous studies have indicated that aging can affect the absorption, tissue distribution, and biological effects of minerals^[5]^.

Urine is not subject to strict homeostatic regulation, allowing it to more directly reflect real-time physiological and pathological changes in the body. Urinary proteomics, as a non-invasive, dynamically monitorable technique enabling continuous sampling and rich in low-molecular-weight proteins and peptides, provides an important tool for investigating physiological and pathological states^[6]^. In particular, post-translational modifications (PTMs)—such as phosphorylation, acetylation, and glycosylation—are widely involved in cellular signal transduction and metabolic regulation, offering a new perspective for deciphering the molecular mechanisms of mineral metabolism^[7]^.

This study selected four clinically common mineral supplements: magnesium L-threonate (MgT), calcium gluconate, zinc gluconate, and polysaccharide-iron complex. MgT is a novel magnesium supplement with high bioavailability, shown to elevate brain magnesium levels, improve neuroplasticity, cross the blood-brain barrier effectively, and exert protective effects on the nervous system^[8]^. Calcium gluconate, an organic calcium salt, is widely used for calcium supplementation due to its good water solubility, high absorption rate, and minimal gastrointestinal irritation compared to inorganic calcium salts^[9]^. Zinc gluconate has demonstrated good intestinal tolerance and immunomodulatory effects in clinical trials, enabling rapid zinc ion release for short-term efficacy^[10]^. The polysaccharide-iron complex is a mild iron supplement with minimal gastrointestinal side effects, high bioavailability, and excellent tolerability, making it suitable for short-term intervention studies^[11]^.

This study aims to administer these four mineral supplements—MgT, calcium gluconate, zinc gluconate, and polysaccharide-iron complex—to Sprague-Dawley (SD) rats of different ages via intragastric gavage over a short term. Using high-throughput proteomic techniques, we systematically analyzed PTM changes in the urine of rats across age groups, exploring the impact of short-term intake of magnesium, calcium, zinc, and iron on protein modifications. The findings provide new clues for understanding the physiological functions of mineral elements and offer a novel perspective for nutritional research.

## 2 Materials and Methods

### 2.1 Experimental Animals and Model Establishment

Data related to the mineral supplement intervention groups and control group rats in this study were derived from previously published research by our laboratory in peer-reviewed journals, as well as findings available on preprint platforms^[12][13][14][15]^. Healthy male Sprague-Dawley (SD) rats (250 ± 20 g) were purchased from Beijing Vital River Laboratory Animal Technology Co., Ltd. All rats were acclimatized for one week under standard conditions: room temperature (22 ± 1)°C, humidity 65%–70%, and a 12 h light/dark cycle. All experimental procedures were reviewed and approved by the Animal Ethics Committee of the College of Life Sciences, Beijing Normal University (Approval No. CLS-AWEC-B-2022-003).

According to the Chinese Dietary Guidelines, the tolerable upper intake levels (UL) for calcium, zinc, and iron are 2000 mg/d, 40 mg/d, and 42 mg/d, respectively. These values were converted to rat-equivalent doses based on body surface area and body weight, resulting in approximate daily intakes of 180 mg/kg·d for calcium, 3.6 mg/kg·d for zinc, and 37.8 mg/kg·d for iron. Corresponding doses for calcium gluconate, zinc gluconate, and polysaccharide-iron complex were approximately 2000 mg/kg·d, 25.3 mg/kg·d, and 28 mg/kg·d (expressed as elemental iron), respectively. Based on previous research by Slutsky et al., the minimal effective dose of magnesium L-threonate (MgT) for enhancing memory function in rats was 604 mg/kg·d (equivalent to 50 mg/kg·d elemental magnesium). MgT is considered the most suitable magnesium salt for oral administration, and a dose of 100 mg/kg·d elemental magnesium was selected^[16][17]^. The daily intakes of mineral supplements and elements for each group are summarized in Table 1.

**Table 1.** Daily intake of mineral supplements and elements in rats.

|  | MgT | Calcium Gluconate | Zinc Gluconate | Polysaccharide-Iron Complex |
| --- | --- | --- | --- | --- |
| Supplement/(mg/kg·d) | 1208 | 3225 | 82 | 60 |
| Element/(mg/kg·d) | 100 | 300 | 11.7 | 28 |

All supplements were dissolved in sterile water to prepare gavage solutions as follows: Zinc gluconate group, 4.2 g zinc gluconate dissolved in 500 mL water; Calcium gluconate group, 16.125 g calcium gluconate dissolved in 500 mL water; Polysaccharide-iron complex group, 3 g complex dissolved in 500 mL water; MgT group, 24.16 g MgT dissolved in 200 mL water.

For the young group, 12 six-week-old rats were randomly divided into four groups (n = 3), each receiving one of the four mineral supplements (MgT, calcium gluconate, zinc gluconate, or polysaccharide-iron complex) via oral gavage. For the adult group, five rats were used and sequentially administered MgT, calcium gluconate, zinc gluconate, or polysaccharide-iron complex at 14, 17, 18, and 19 weeks of age, with washout periods between treatments. All rats were gavaged once daily for four consecutive days. Urine samples were collected one day before gavage as the control group and on the fourth day of gavage as the experimental group for self-paired comparison. All collected urine samples were temporarily stored at -80°C.

### 2.2 Urine Sample Processing

A 2 mL aliquot of urine was thawed and centrifuged at 12,000 × g for 30 min at 4°C. The supernatant was collected, and 40 μL of 1M dithiothreitol (DTT, Sigma) stock solution was added to achieve a final concentration of 20 mM. The mixture was thoroughly vortexed, incubated at 37°C in a metal bath for 60 min, and then cooled to room temperature.

Subsequently, 100 μL of iodoacetamide (IAA, Sigma) stock solution was added to achieve the desired IAM concentration. After thorough mixing, the reaction was carried out at room temperature in the dark for 45 min. The sample was then transferred to a new centrifuge tube, mixed with three volumes of pre-chilled absolute ethanol, and stored at -20°C for 24 h to precipitate proteins. After precipitation, the sample was centrifuged at 10,000 × g for 30 min at 4°C, and the supernatant was discarded. The protein pellet was dried and redissolved in 200 μL of 20 mM Tris solution. After another centrifugation step, the supernatant was retained, and the protein concentration was determined using the Bradford method.

The filter-aided sample preparation (FASP) method was employed for further processing. The urinary protein extract was loaded onto a 10 kDa ultrafiltration tube (Pall, Port Washington, NY, United States) and washed three times with 20 mM Tris solution. Proteins were redissolved in 30 mM Tris solution, and trypsin (Trypsin Gold, Promega, Fitchburg, WI, United States) was added at a ratio of 1:50 (enzyme-to-protein) for digestion at 37°C for 16 h.

The digested peptides were collected as the peptide mixture. After desalting using an Oasis HLB solid-phase extraction column, the peptides were vacuum-dried and stored at -80°C. The lyophilized peptide powder was redissolved in 30 μL of 0.1% formic acid solution, and the peptide concentration was determined using a BCA kit. The peptide concentration was adjusted to 0.5 μg/μL, and 4 μL from each sample was pooled to prepare a mixed sample.

### 2.3 LC-MS/MS Analysis

Peptides were redissolved in 0.1% formic acid solution and diluted to a concentration of 0.5 μg/μL. Separation was performed using a Thermo Easy-nLC 1200 nano-liquid chromatography system under the following conditions: a 90-minute elution time with mobile phase A (0.1% formic acid in water) and mobile phase B (80% acetonitrile), using a gradient elution program. The separated peptides were analyzed using an Orbitrap Fusion Lumos Tribrid mass spectrometer in data-independent acquisition (DIA) mode.

### 2.4 Open-pSearch for Unrestricted Modification Identification

The pFind Studio software (version 3.2.1, Institute of Computing Technology, Chinese Academy of Sciences) was used to perform unrestricted modification searches on the raw mass spectrometry data of each sample, with default parameters applied during the search. The database used was the *Rattus norvegicus* protein database downloaded from UniProt (https://www.uniprot.org) in September 2024. The instrument type was set to HCD-FTMS, trypsin was selected as the enzyme with full specificity, and a maximum of two missed cleavages were allowed. The mass error tolerance for both precursor and fragment ions was set to ±20 ppm, and the open search mode was selected. The false discovery rate (FDR) threshold at the peptide level was set to 1%.

### 2.5 Bioinformatic Analysis of Protein Post-Translational Modifications

After completing the unrestricted modification search, the modification identification results (PROTEIN files) for each sample were obtained. A Python script named *pFind_protein_contrast_script* was downloaded from GitHub (https://github.com/daheitu/scripts_for_pFind3_protocol.io) to aggregate modification identification results across samples. Differential modified proteins between experimental and control groups were screened based on the following criteria: fold change (FC) ≥ 2.0 or ≤ 0.5, and a two-tailed paired Student’s t-test p-value < 0.01. Proteins with differential modifications were annotated and functionally characterized using the UniProt database. Relevant literature was retrieved from PubMed (https://pubmed.ncbi.nlm.nih.gov) to further analyze the potential functions of these modifications.

## 3 Results Analysis

### 3.1 Effects of Magnesium L-Threonate (MgT) Intervention on Post-Translational Modifications in the Rat Urinary Proteome

#### 3.1.1 Analysis of PTMs in the Urinary Proteome of 9-Week-Old Rats after MgT Intervention

A comparison of post-translational modifications between the experimental group (after 4 days of gavage) and the control group (before gavage) revealed that one differential PTM was identified in the 9-week-old group, corresponding to one protein. Details are listed in Table 2.

**Table 2.** Differential PTMs screened with FC ≥ 2.0 or ≤ 0.5 and P < 0.01 in the MgT 9-week-old group.

| Uniprot ID | Peptide | Modification | Control group | Experimental group | Fold change | P value |
| --- | --- | --- | --- | --- | --- | --- |
| P12346 | EGVCPEGSIDSAPVK | 4,Carbamidomethyl[C]; | 5.0 | 14.3 | 2.9 | 5.06E-03 |

A literature search was conducted in the PubMed database for the protein corresponding to the identified PTM.

P12346, Serotransferrin (FC = 2.9, P = 5.06E-03), is primarily responsible for iron transport and cellular uptake. Transferrin saturation (TSAT) is an internationally recognized biomarker of iron nutritional status. TSAT decreases during iron deficiency, leading to compensatory increases in serotransferrin synthesis, while it elevates during iron overload. Currently, no studies have reported a direct relationship between serotransferrin and changes in magnesium levels in the body^[18]^.

#### 3.1.2 Analysis of PTMs in the Urinary Proteome of 14-Week-Old Rats after MgT Intervention

Comparison of post-translational modifications between the experimental group (after 4 days of gavage) and the control group (before gavage) showed that 29 differential PTMs were identified in the 14-week-old group, involving 19 proteins. Detailed information is presented in Table 3.

**Table 3.**
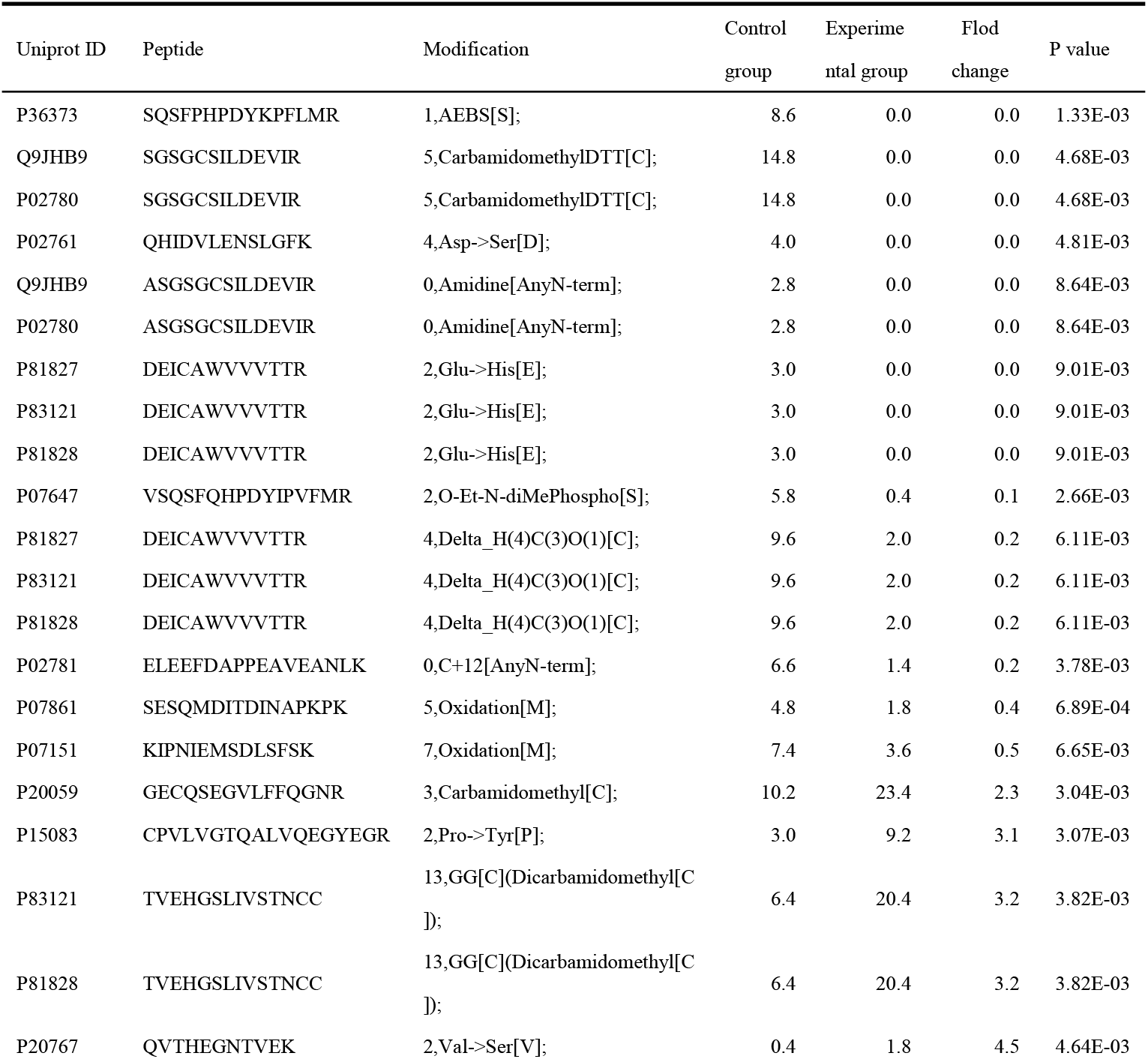
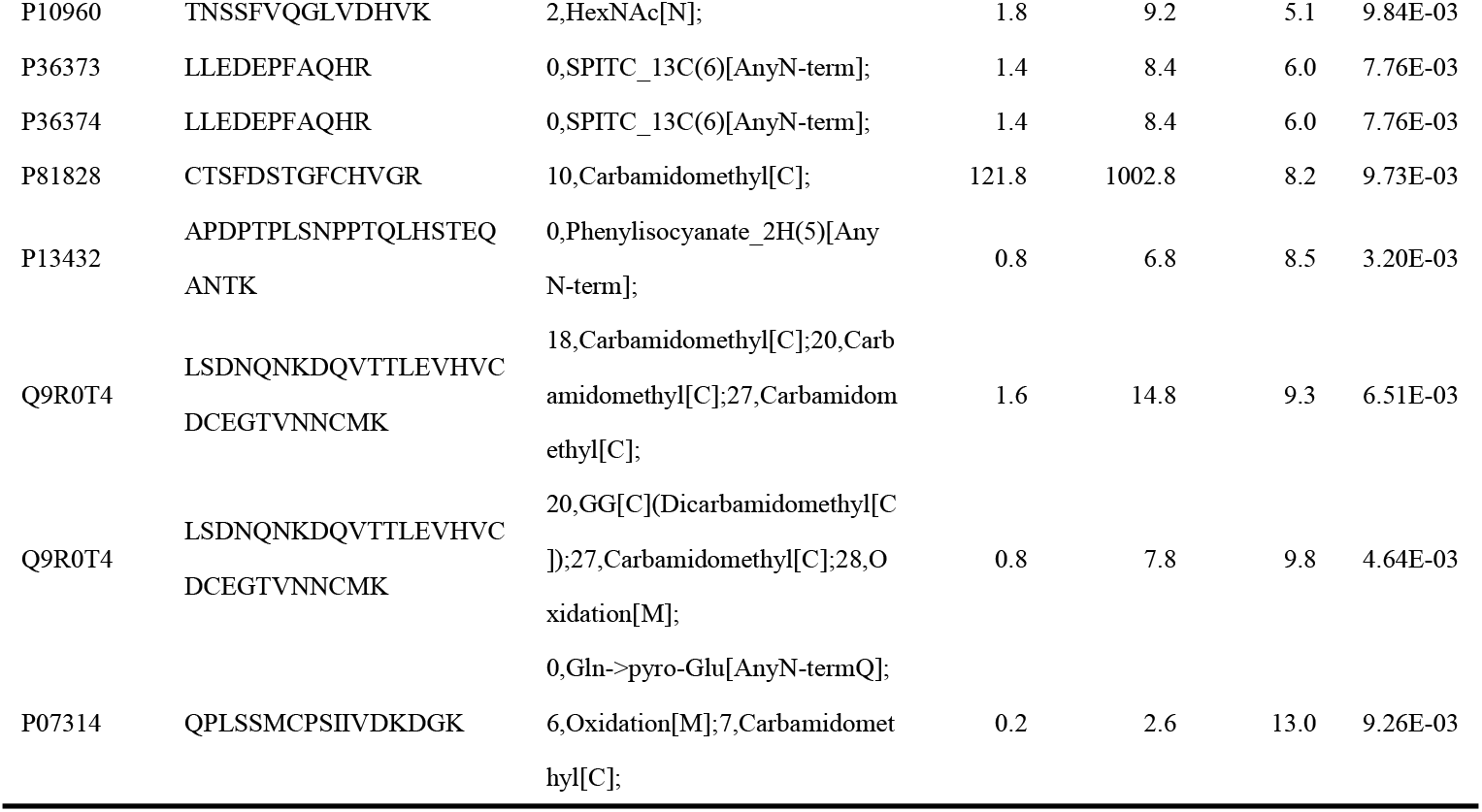
Differential PTMs screened with FC ≥ 2.0 or ≤ 0.5 and P < 0.01 in the MgT 14-week-old group.

A literature search was conducted in the PubMed database for proteins carrying the identified post-translational modifications. Some of these proteins have been reported in previous studies to be associated with changes in magnesium concentration in the body.

P07861, Neprilysin (FC=0.4, P=6.89E-04). Changes in magnesium concentration may affect the activity of Neprilysin. Specifically, low magnesium concentrations (0.0 and 0.4 mM) led to a 50% decrease in Neprilysin activity, while the protein level of Neprilysin remained unchanged^[19]^. This suggests that changes in magnesium concentration may regulate its function by affecting Neprilysin’s activity.

P07151, Beta-2-microglobulin (FC=0.5, P=6.65E-03). Studies have found that Mg is a significant independent factor for increased serum concentration of Beta-2-microglobulin, and the level of this protein was negatively correlated with serum magnesium concentration^[20]^.

### 3.2 Effects of Calcium Gluconate Intervention on Post-Translational Modifications in the Rat Urinary Proteome

#### 3.2.1 Analysis of PTMs in the Urinary Proteome of 9-Week-Old Rats after Calcium Gluconate Intervention

Comparison of post-translational modifications between the experimental group (after 4 days of gavage) and the control group (before gavage) revealed that one differential PTM was identified in the 9-week-old group, corresponding to one protein. Detailed information is presented in Table 4.

**Table 4.**
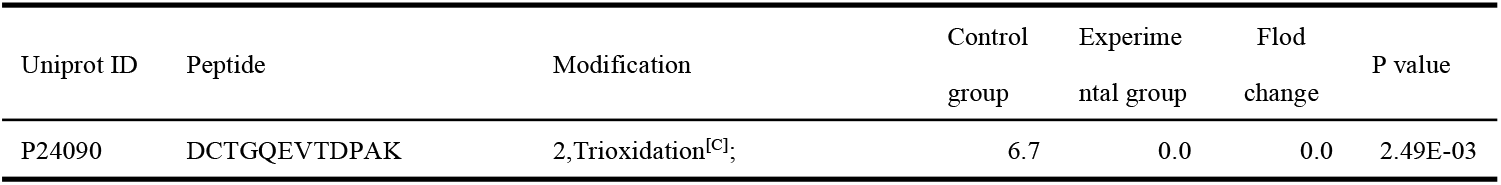
Differential PTMs screened with FC ≥ 2.0 or ≤ 0.5 and P < 0.01 in the Calcium Gluconate 9-week-old group.

| Uniprot ID | Peptide | Modification | Control group | Experimental group | Flod change | P value |
| --- | --- | --- | --- | --- | --- | --- |
| P24090 | DCTGQEVTDPAK | 2,Trioxidation<sup>[C]</sup>; | 6.7 | 0.0 | 0.0 | 2.49E-03 |

A literature search was performed in the PubMed database for proteins carrying the identified post-translational modifications.

P24090, Alpha-2-HS-glycoprotein (FC=0, P=2.49E-03). Studies have found that calcification load is negatively correlated with serum Alpha-2-HS-glycoprotein levels. This protein can form colloidal calcio-protein particles that regulate calcium solubility and bioavailability. These particles circulate in the bloodstream and contribute to the maintenance of calcium homeostasis^[21]^.

#### 3.2.2 Analysis of PTMs in the Urinary Proteome of 16-Week-Old Rats after Calcium Gluconate Intervention

Comparison of post-translational modifications between the experimental group (after 4 days of gavage) and the control group (before gavage) revealed that 11 differential PTMs were identified in the 16-week-old group, involving 10 proteins. Detailed information is presented in Table 5.

**Table 5.**
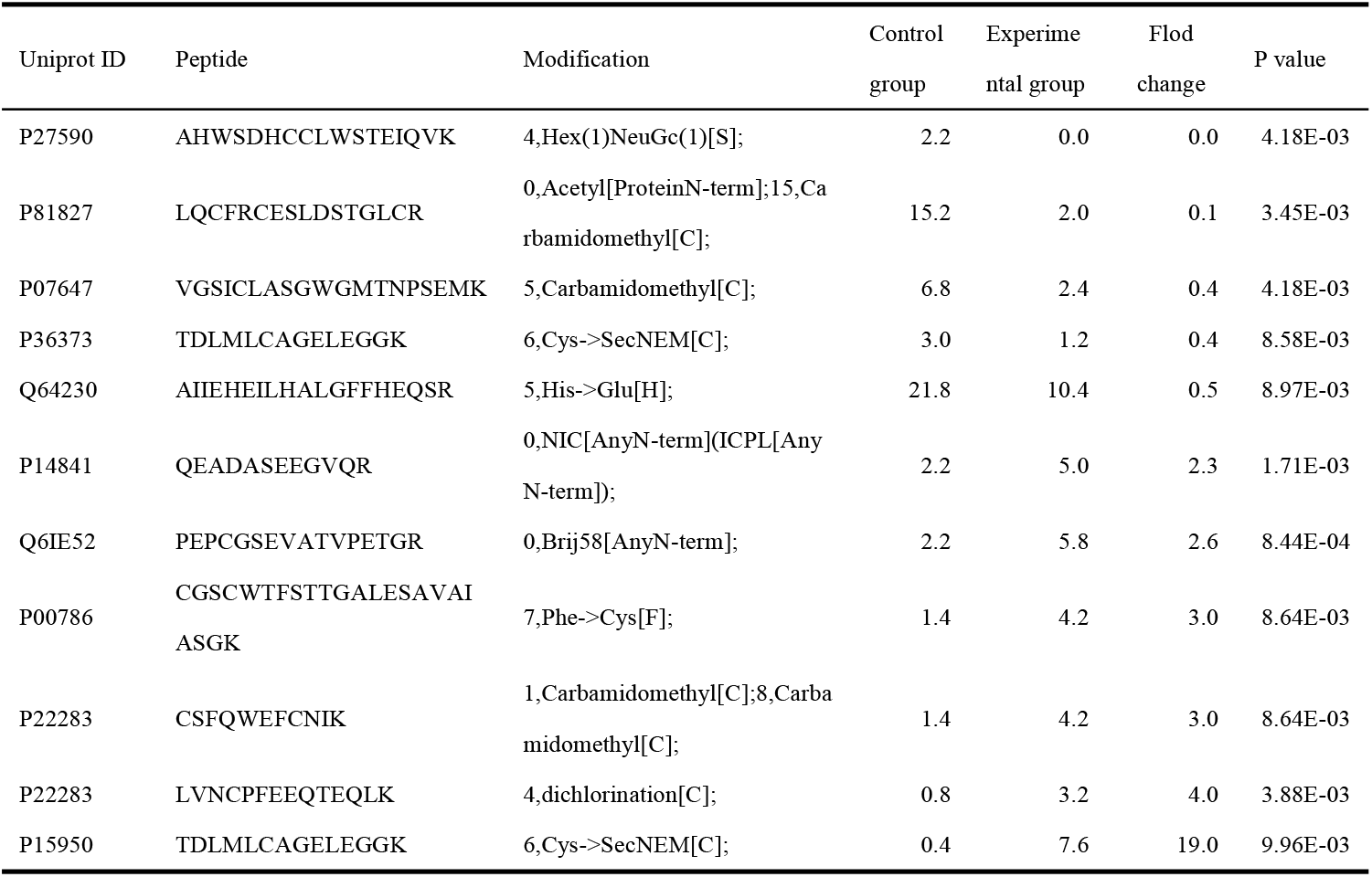
Differential PTMs screened with FC ≥ 2.0 or ≤ 0.5 and P < 0.01 in the Calcium Gluconate 16-week-old group.

A literature search was conducted in the PubMed database for proteins carrying the identified post-translational modifications. Some of these proteins have been reported in previous studies to be associated with changes in calcium concentration in the body.

P27590, Uromodulin (FC=0, P=4.18E-03), is a secretory protein primarily expressed in the thick ascending limb of the loop of Henle in the kidney. It participates in urine concentration and calcium metabolism, regulating systemic calcium balance. In calcium metabolism regulation, uromodulin significantly stimulates the expression of the transient receptor potential vanilloid 5 (TRPV5) on the apical membrane of renal tubular epithelial cells by inhibiting endocytosis, thereby enhancing calcium reabsorption efficiency. This regulatory mechanism serves as an important protective strategy against urinary stone formation^[22]^.

Q6IE52, Murinoglobulin-2 (FC=2.6, P=8.44E-04). Studies have shown that β2-microglobulin, which is homologous to Murinoglobulin-2, binds to calcium at physiological pH, leading to conformational changes and precipitation of the protein into amorphous forms, which subsequently transform into amyloid aggregates. These aggregates exist as microscopic particles and do not progress into larger amyloid formations. However, when renal function is impaired, particularly during dialysis, the concentration of β2-microglobulin may temporarily increase, resulting in large aggregates that deposit in bones and joints and transform into amyloid during dialysis-related amyloidosis^[23]^.

### 3.3 Effects of Zinc Gluconate Intervention on Post-Translational Modifications in the Rat Urinary Proteome

#### 3.3.1 Analysis of PTMs in the Urinary Proteome of 9-Week-Old Rats after Zinc Gluconate Intervention

Comparison of post-translational modifications between the experimental group (after 4 days of gavage) and the control group (before gavage) revealed that 6 differential PTMs were identified in the 9-week-old group, involving 6 proteins. Detailed information is presented in Table 6.

**Table 6.** Differential PTMs screened with FC ≥ 2.0 or ≤ 0.5 and P < 0.01 in the Zinc Gluconate 9-week-old group.

| Uniprot ID | Peptide | Modification | Control group | Experimental group | Flod change | P value |
| --- | --- | --- | --- | --- | --- | --- |
| Q63041 | CGNKVAEVPALVQK | 0,Ethylphosphate[AnyN-term]; | 3.3 | 0.0 | 0.0 | 9.85E-03 |
| P81827 | QTYPDEICAWVVVTTR | 0,Succinyl_2H(4)[AnyN-term]; | 26.0 | 0.3 | 0.0 | 8.66E-03 |
| P83121 | QTYPDEICAWVVVTTR | 0,Succinyl_2H(4)[AnyN-term]; | 26.0 | 0.3 | 0.0 | 8.66E-03 |
| P81828 | QTYPDEICAWVVVTTR | 0,Succinyl_2H(4)[AnyN-term]; | 26.0 | 0.3 | 0.0 | 8.66E-03 |
| P02761 | VFMQHIDVLENSLGFK | 0,Cytopiloyne[AnyN-term]; | 3.7 | 0.3 | 0.1 | 9.85E-03 |
| P12346 | DCTGNFCLFR | 2,Carbamidomethyl[C];7,Carbamidomethyl[C]; | 20.3 | 8.3 | 0.4 | 2.31E-03 |

A literature search was conducted in the PubMed database for proteins carrying the identified post-translational modifications. Some of these proteins have been reported in previous studies to be associated with changes in zinc concentration in the body.

P12346, Serotransferrin (FC=0.4, P=2.31E-03), is one of the important zinc-binding proteins in plasma. It is closely related to dietary zinc intake and zinc homeostasis in the body, and participates in regulating zinc absorption and utilization, as well as zinc transport and tissue distribution^[24]^.

#### 3.3.2 Analysis of PTMs in the Urinary Proteome of 17-Week-Old Rats after Zinc Gluconate Intervention

Comparison of post-translational modifications between the experimental group (after 4 days of gavage) and the control group (before gavage) revealed that 14 differential PTMs were identified in the 17-week-old group, involving 11 proteins. Detailed information is presented in Table 7.

**Table 7.**
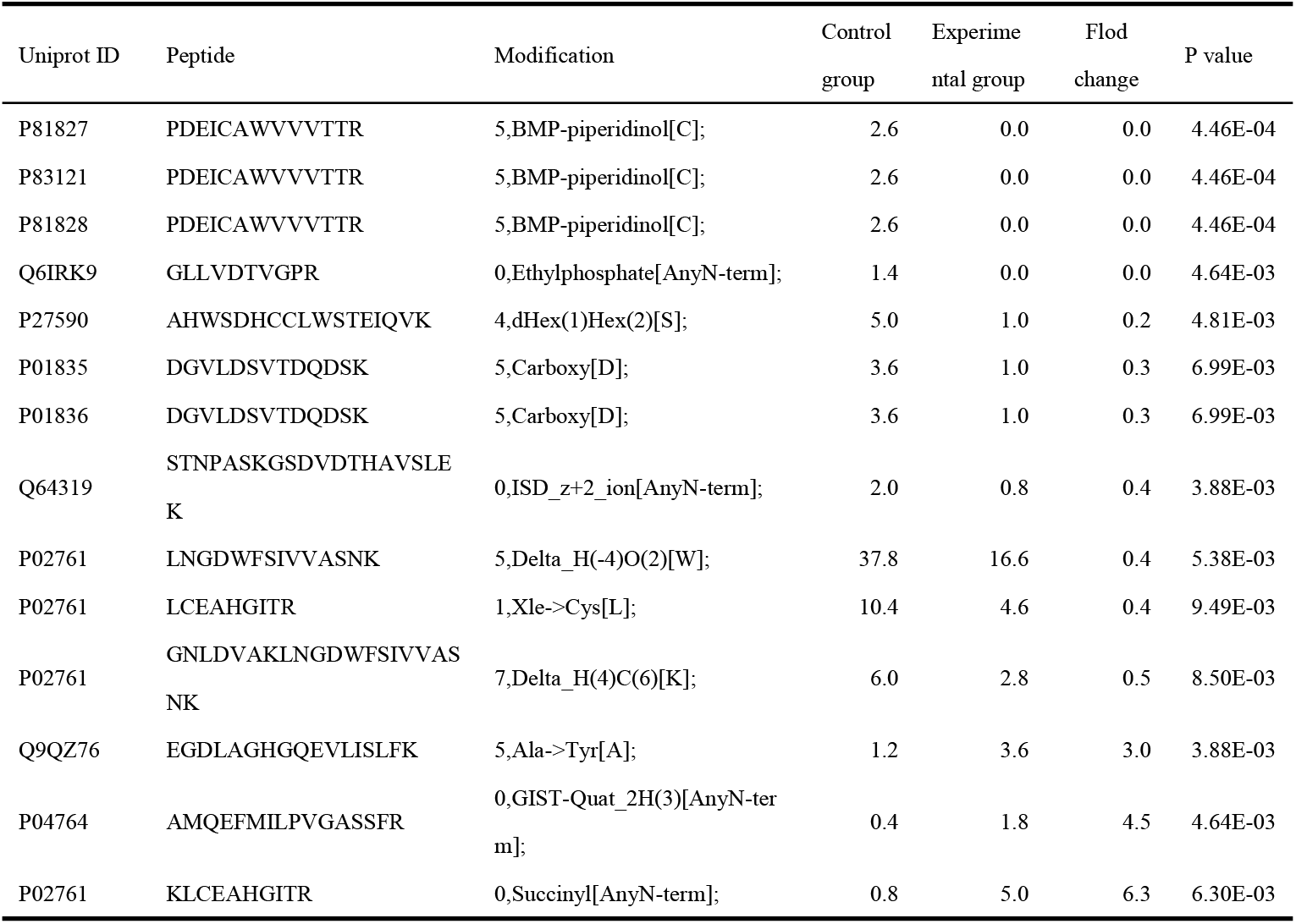
Differential PTMs screened with FC ≥ 2.0 or ≤ 0.5 and P < 0.01 in the Zinc Gluconate 17-week-old group.

A literature search was conducted in the PubMed database for proteins carrying the identified post-translational modifications. Some of these proteins have been reported in previous studies to be associated with changes in zinc concentration in the body.

Q64319, Amino acid transporter heavy chain SLC3A1 (FC=0.4, P=3.88E-03). SLC3A1, SLC30, and SLC39 are different members of the solute carrier (SLC) family. While they have distinct functions, potential interconnections exist among them. The SLC30 family primarily functions in zinc efflux from cells or zinc sequestration within specific cytoplasmic compartments when intracellular zinc levels rise, thereby maintaining physiological zinc concentrations^[25]^. Conversely, the SLC39 family is mainly responsible for zinc uptake into cells, ensuring dynamic balance of intracellular zinc ions^[26]^.The known function of SLC3A1 primarily focuses on amino acid transport. Although no direct evidence currently links SLC3A1 to zinc ions, considering the complex interactions among SLC superfamily members within cellular metabolic networks and the potential indirect relationship between amino acid metabolism and zinc homeostasis, the possibility of its association with zinc regulation warrants further investigation.

P01836, Ig kappa chain C region, A allele (FC=0.3, P=6.99E-03). Zinc is an essential trace element for normal immune system function, participating in the regulation of immune cell activity and function. Both deficiency and excess of zinc can affect immune responses, including antibody production and function^[27]^

### 3.4 Effects of Polysaccharide-Iron Complex Intervention on Post-Translational Modifications in the Rat Urinary Proteome

#### 3.4.1 Analysis of PTMs in the Urinary Proteome of 9-Week-Old Rats after Polysaccharide-Iron Complex Intervention

Comparison of post-translational modifications between the experimental group (after 4 days of gavage) and the control group (before gavage) revealed that 38 differential PTMs were identified in the 9-week-old group, involving 31 proteins. Detailed information is presented in Table 8.

**Table 8.**
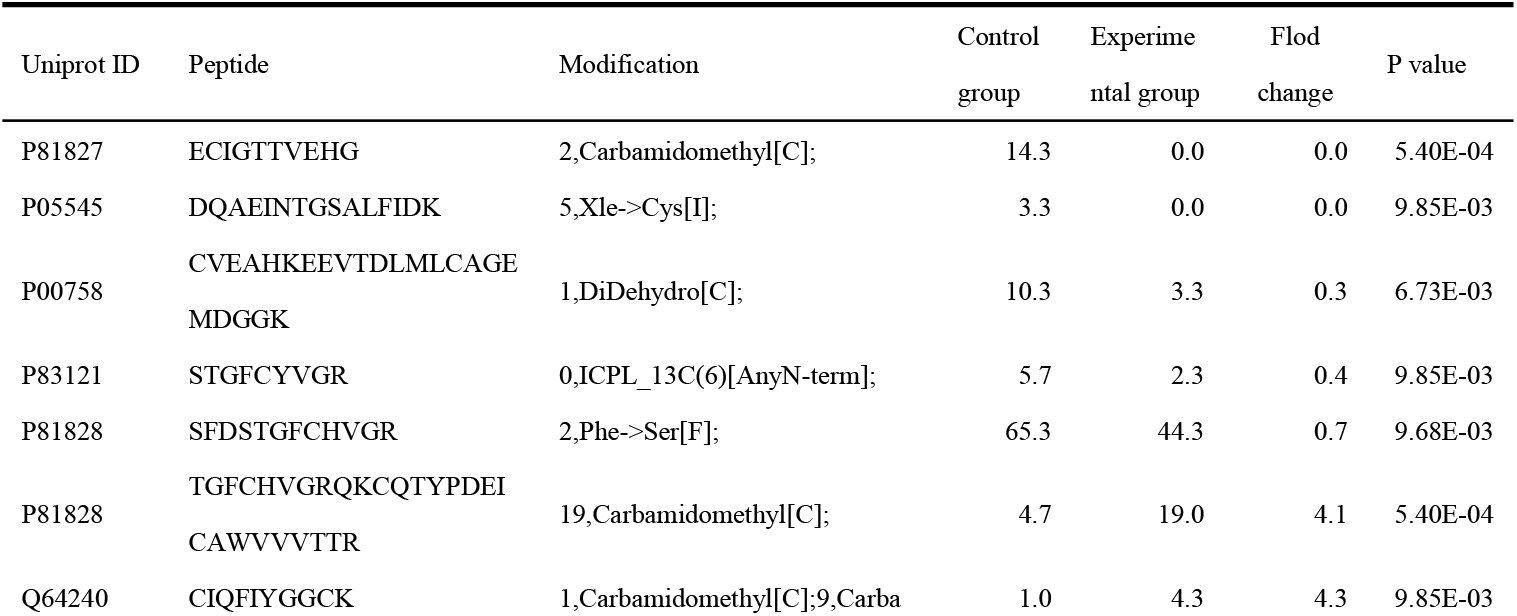
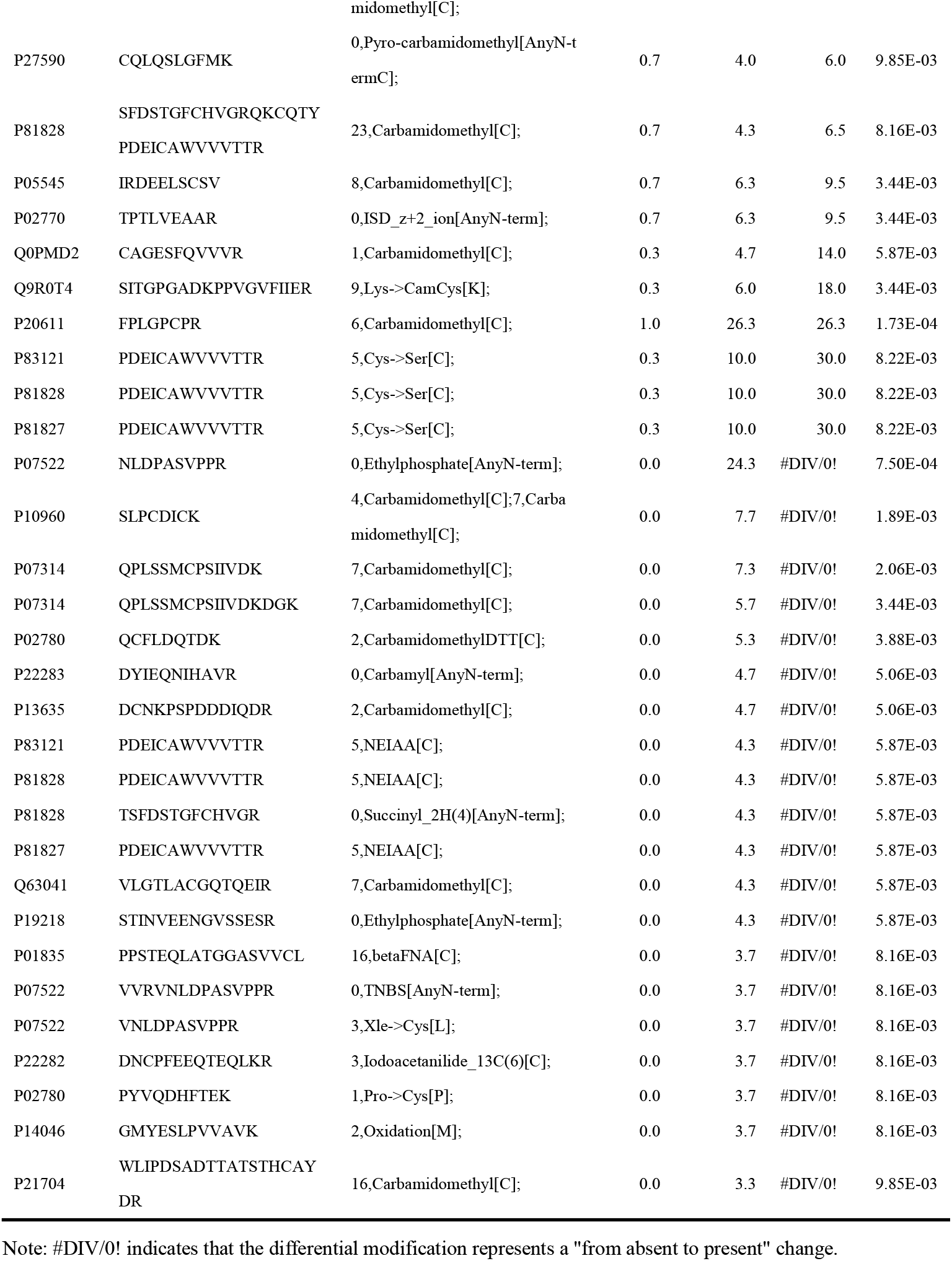
Differential PTMs screened with FC ≥ 2.0 or ≤ 0.5 and P < 0.01 in the Polysaccharide-Iron Complex 9-week-old group.

A literature search was conducted in the PubMed database for proteins carrying the identified post-translational modifications. Some of these proteins have been reported in previous studies to be associated with changes in iron concentration in the body.

P02770, Albumin (FC=9.5, P=3.44E-03). Studies have shown that serum albumin levels are significantly lower in patients with iron deficiency anemia compared to healthy controls, and albumin levels are positively correlated with serum iron, transferrin saturation, and ferritin levels^[28]^. This suggests that changes in iron concentration may affect albumin levels.

Q9R0T4, Cadherin-1 (FC=18, P=3.44E-03). Research has found that Cadherin-1 expression is significantly decreased in iron-overloaded hepatocytes, leading to impaired intercellular adhesion function^[29]^. P20611, Lysosomal acid phosphatase (FC=26.3, P=1.73E-04). In iron-overloaded hepatocytes, both the number of lysosomes and the activity of lysosomal acid phosphatase are increased^[30]^.

Among proteins showing “from absent to present” differential modifications: P13635, Ceruloplasmin (P=5.06E-03). Under increased iron load, ceruloplasmin promotes iron loading onto transferrin by enhancing its oxidase activity, thereby accelerating iron clearance. Its activity may be dynamically adjusted according to dietary iron load^[31]^.Q63041, Alpha-1-macroglobulin (P=5.87E-03). This protein can bind free heme with high affinity. Its plasma concentration and urinary excretion significantly increase during dietary iron overload or hemolysis-induced heme release, suggesting its potential as a biomarker for iron overload or hemolytic dietary stress^[32]^.P21704, Deoxyribonuclease-1 (P=9.85E-03). The activity of this protein is directly related to dietary iron intake. High-iron diets can lead to iron overload in the body, subsequently causing increased reactive oxygen species and apoptosis, ultimately resulting in elevated DNase-1 activity^[33]^.

#### 3.4.2 Analysis of PTMs in the Urinary Proteome of 16-Week-Old Rats after Polysaccharide-Iron Complex Intervention

Comparison of post-translational modifications between the experimental group (after 4 days of gavage) and the control group (before gavage) revealed that 61 differential PTMs were identified in the 17-week-old group, involving 34 proteins. Detailed information is presented in Table 9.

**Table 9.**
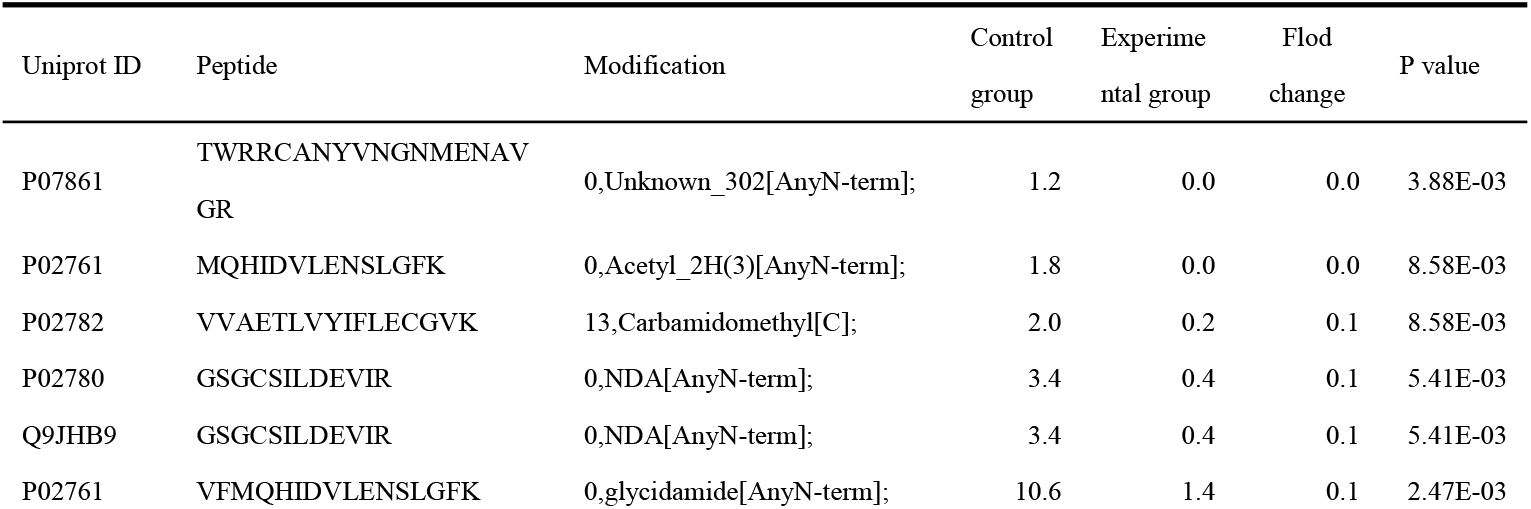
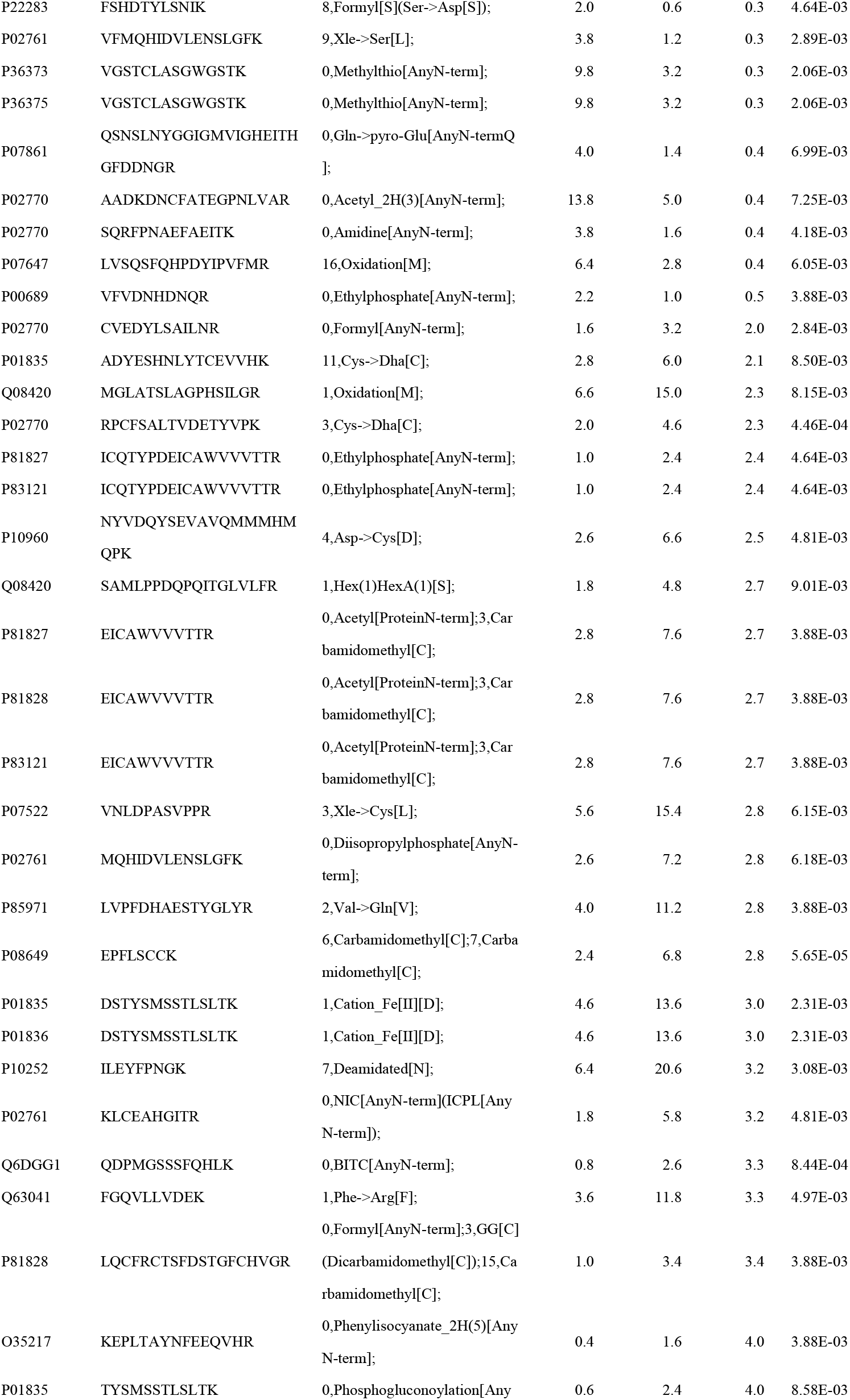
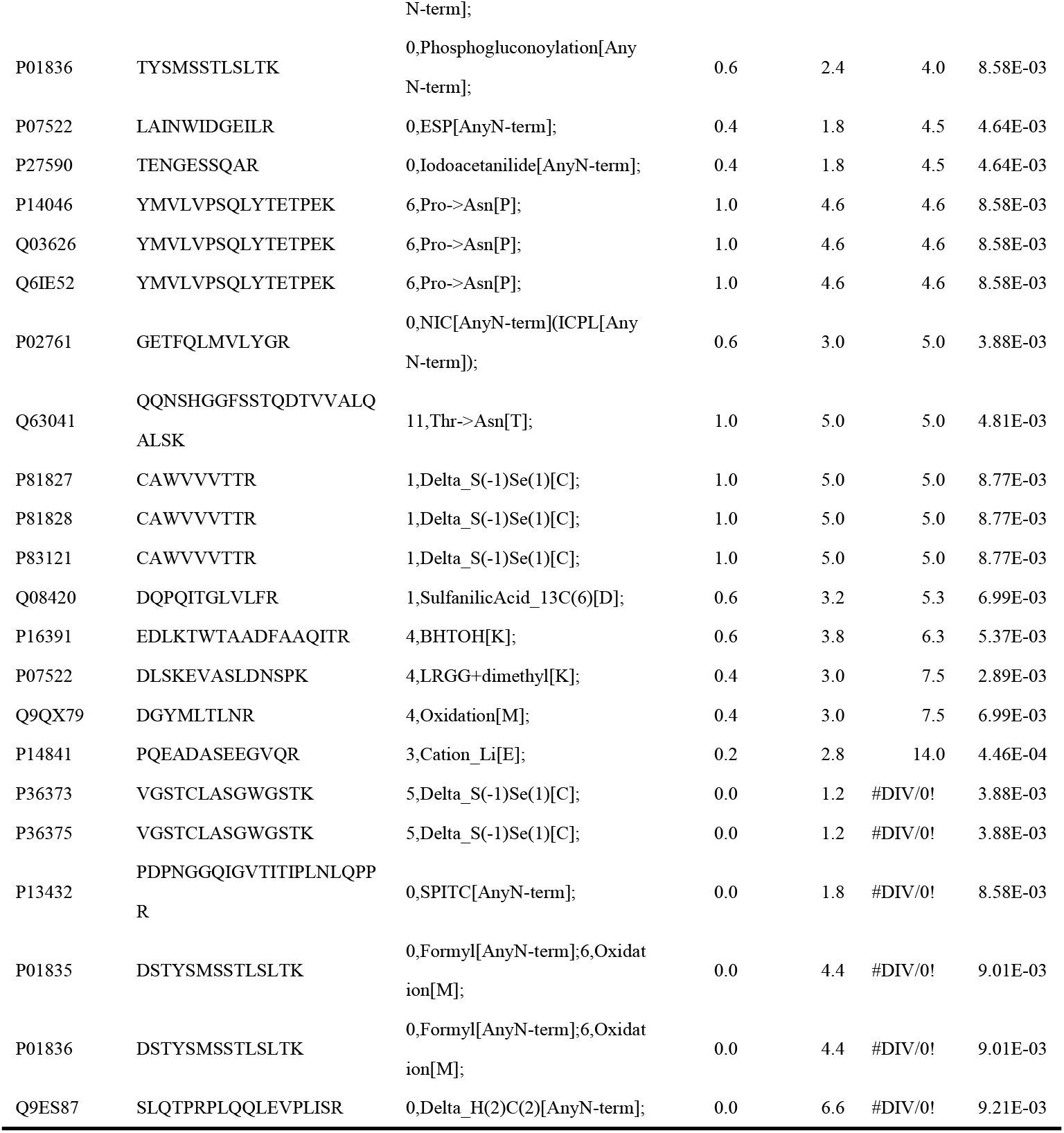
Differential PTMs screened with FC ≥ 2.0 or ≤ 0.5 and P < 0.01 in the Polysaccharide-Iron Complex 17-week-old group.

A literature search was conducted in the PubMed database for proteins carrying the identified post-translational modifications. Some of these proteins have been reported in previous studies to be associated with changes in iron concentration in the body.

The 17-week-old group shared several iron metabolism-related differentially modified proteins with the 9-week-old group, including Albumin and Alpha-1-macroglobulin. Additionally:

P02782, Prostatic steroid-binding protein C1 (FC=0.1, P=8.58E-03). This protein is associated with hepcidin synthesis and secretion in prostate cancer cells. Hepcidin, a key iron-regulatory protein, shows significantly increased expression and secretion in prostate cancer cells and tissues^[34]^. PSBP C1 may influence iron metabolism by regulating hepcidin expression. P00689, Pancreatic alpha-amylase (FC=0.5, P=3.88E-03). Studies have found significant correlations between iron regulatory proteins and pancreatic α-amylase levels. The regulation of iron metabolism may affect pancreatic cell physiology, thereby influencing the synthesis and secretion of α-amylase^[35]^.

Notably, a common differential modification was identified in both the 9-week and 17-week groups: the modification “3,Xle->Cys[L]” on the “VNLDPASVPPR” sequence of Pro-epidermal growth factor (Pro-EGF), with FC values of #DIV/0! (indicating “from absent to present” change) and 2.8, respectively. The epidermal growth factor receptor (EGFR) plays a crucial role in iron metabolism by regulating the distribution of transferrin receptor 1^[36]^. As a member of the EGF family, the expression and function of Pro-EGF may be regulated by the EGFR signaling pathway, thereby indirectly influencing iron metabolism.

## 4 Discussion

This study investigated alterations in post-translational modifications (PTMs) of the urinary proteome in rats following short-term intragastric administration of four common mineral supplements: magnesium L-threonate (MgT), calcium gluconate, zinc gluconate, and polysaccharide-iron complex. Changes in protein concentration and PTMs represent complementary information, capturing the effects of subtle, short-term interventions from two distinct dimensions.

At the PTM level, all mineral intervention groups exhibited differential modification sites, with the polysaccharide-iron complex group showing the most pronounced changes, suggesting that iron metabolism may exert a stronger influence on protein modification regulation. Notably, the proteins carrying these differential modifications are involved in diverse biological functions, including metal ion binding, immune regulation, and oxidative stress response. These findings not only align with existing knowledge of mineral-protein interactions but also highlight urinary PTM analysis as a promising tool for elucidating the physiological functions of minerals. Age-related differences were also observed in this study. The same mineral supplement induced significantly different PTM changes in rats of different ages, consistent with known age-dependent variations in absorption efficiency, enzyme activity, and hormonal environments^[5]^.This further emphasizes the importance of considering age as a critical factor in nutritional interventions.

However, it should be noted that the functional annotations of most identified PTMs in mineral metabolism remain incomplete, which limits the depth of analysis for specific modification events in this study. Future research should combine targeted proteomics with functional experiments to further clarify the roles of key modification sites in mineral homeostasis and related physiological functions. Additionally, this study has several limitations. While we focused on the correlation between differentially modified proteins and the metabolism of magnesium, calcium, zinc, and iron, and although the supplements used are widely applied in clinical and basic research with minimal non-specific effects from carrier components, other constituents such as L-threonate anions, gluconate, and polysaccharide carriers may still potentially influence the results. Furthermore, the relatively small sample size may affect the generalizability and statistical power of the findings. Nevertheless, as an exploratory study, its primary aim was to preliminarily characterize the features and trends of mineral-protein modification interactions and provide clues and directions for subsequent mechanistic investigations. Future studies could consider expanding the sample size, including appropriate vehicle control groups, and employing targeted validation, functional experiments, and multi-omics integration strategies to further explore key modification events regulated by minerals and their physiological and pathological significance, thereby providing a theoretical basis for precision nutrition interventions.

## 5 Conclusion

In summary, this study preliminarily explored changes in the rat urinary proteome following short-term supplementation with magnesium L-threonate (MgT), calcium gluconate, zinc gluconate, and polysaccharide-iron complex from a PTM perspective. The findings revealed that some proteins with differential modifications were consistent with those reported in previous studies on mineral metabolism, providing new clues for understanding the metabolic processes and biological functions of magnesium, calcium, zinc, and iron in rats. This work opens a new window for nutritional research.

## References

[1] de Baaij JH, Hoenderop JG, Bindels RJ. Magnesium in man: implications for health and disease. Physiol Rev. 2015 Jan;95(1):1–46.

[2] Peacock M. Calcium metabolism in health and disease. Clin J Am Soc Nephrol. 2010 Jan;5 Suppl 1:S23–30. doi: 10.2215/CJN.05910809. PMID: 20089499.

[3] Hojyo S, Fukada T. Roles of Zinc Signaling in the Immune System. J Immunol Res. 2016;2016:6762343. doi: 10.1155/2016/6762343. Epub 2016 Oct 31. PMID: 27872866; PMCID: PMC5107842.

[4] Coates TD. Physiology and pathophysiology of iron in hemoglobin-associated diseases. Free Radic Biol Med. 2014 Jul;72:23–40. doi: 10.1016/j.freeradbiomed.2014.03.039. Epub 2014 Apr 12. PMID: 24726864; PMCID: PMC4940047.

[5] Matsui Y, Takemura M, Harada A, Ando F, Shimokata H. Divergent Significance of Bone Mineral Density Changes in Aging Depending on Sites and Sex Revealed through Separate Analyses of Bone Mineral Content and Area. J Osteoporos. 2012;2012:642486. doi: 10.1155/2012/642486. Epub 2012 Nov 25. PMID: 23227425; PMCID: PMC3512306.

[6] Wei J, Gao Y. Early disease biomarkers can be found using animal models urine proteomics. Expert Rev Proteomics. 2021 May;18(5):363–378. doi: 10.1080/14789450.2021.1937133. Epub 2021 Jun 7. PMID: 34058951.

[7] D’Amore C, Salvi M. Editorial of Special Issue “Protein Post-Translational Modifications in Signal Transduction and Diseases”. Int J Mol Sci. 2021 Feb 24;22(5):2232. doi: 10.3390/ijms22052232. PMID: 33668127; PMCID: PMC7956322.

[8] Fu C, Huang L, Lian C, Yue J, Lin P, Xu L, Lai W, Gao C, Li C, Long Y. Effects of long-term magnesium L-threonate supplementation on neuroinflammation, demyelination and blood-brain barrier integrity in mice with neuromyelitis optica spectrum disorder. Brain Res. 2025 Jan 1;1846:149234. doi: 10.1016/j.brainres.2024.149234. Epub 2024 Sep 10. PMID: 39260790.

[9] Watanabe DHM, Doelman J, Steele MA, Guan LL, Seymour DJ, Penner GB. A comparison of post-ruminal provision of Ca-gluconate and Ca-butyrate on growth performance, gastrointestinal barrier function, short-chain fatty acid absorption, intestinal histology, and brush-border enzyme activity in beef heifers. J Anim Sci. 2023 Jan 3;101:skad050. doi: 10.1093/jas/skad050. PMID: 36799118; PMCID: PMC10022388.

[10] Wang Y, Xiao J, Wei S, Su Y, Yang X, Su S, Lan L, Chen X, Huang T, Shan Q. Protective effect of zinc gluconate on intestinal mucosal barrier injury in antibiotics and LPS-induced mice. Front Microbiol. 2024 May 23;15:1407091. doi: 10.3389/fmicb.2024.1407091. PMID: 38855764; PMCID: PMC11157515.

[11] Yan X, Zhang Q, Wang T, Luo Y, Sha X. Evaluation of Different Polysaccharide-Iron Complex Preparations In Vitro and In Vivo. Pharmaceutics. 2025 Feb 23;17(3):292. doi: 10.3390/pharmaceutics17030292. PMID: 40142956; PMCID: PMC11945278.

[12] Shen Z, Yang M, Wang H, Liu Y, Gao Y. Changes in the urinary proteome of rats after short-term intake of magnesium L-threonate(MgT). Front Nutr. 2023 Dec 21;10:1305738. doi: 10.3389/fnut.2023.1305738. PMID: 38188875; PMCID: PMC10768015.

[13] Shen, Ziyun, Minhui Yang, Haitong Wang, and Youhe Gao. “Changes of urinary proteome in rats after intragastric administration of zinc gluconate.” *bioRxiv* 4 Mar. 2024. 10.1101/2024.03.04.583149

[14] Shen, Ziyun, Minhui Yang, Haitong Wang, and Youhe Gao. “Changes of urinary proteome in rats after intragastric administration of calcium gluconate.” *bioRxiv* 4 Mar. 2024. 10.1101/2024.03.04.583150

[15] Shen, Ziyun, Minhui Yang, Haitong Wang, and Youhe Gao. “Changes of urine proteome after intragastric administration of polysaccharide iron complex in rats.” *bioRxiv* 5 Mar. 2024. 10.1101/2024.03.05.583147

[16] Slutsky I, Abumaria N, Wu LJ, Huang C, Zhang L, Li B, Zhao X, Govindarajan A, Zhao MG, Zhuo M, Tonegawa S, Liu G. Enhancement of learning and memory by elevating brain magnesium. Neuron. 2010 Jan 28;65(2):165–77. doi: 10.1016/j.neuron.2009.12.026. PMID: 20152124.

[17] Sadir S, Tabassum S, Emad S, Liaquat L, Batool Z, Madiha S, Shehzad S, Sajid I, Haider S. Neurobehavioral and biochemical effects of magnesium chloride (MgCl2), magnesium sulphate (MgSO4) and magnesium-L-threonate (MgT) supplementation in rats: A dose dependent comparative study. Pak J Pharm Sci. 2019 Jan;32(1(Supplementary)):277–283. PMID: 30829204.

[18] Pfeiffer CM, Looker AC. Laboratory methodologies for indicators of iron status: strengths, limitations, and analytical challenges. Am J Clin Nutr. 2017 Dec;106(Suppl 6):1606S–1614S. doi: 10.3945/ajcn.117.155887. Epub 2017 Oct 25. PMID: 29070545; PMCID: PMC5701713.

[19] Chui D, Chen Z, Yu J, et al.; Liang Zhou. Magnesium in Alzheimer’s disease. In: Vink R, Nechifor M, editors. Magnesium in the Central Nervous System [Internet]. Adelaide (AU): University of Adelaide Press; 2011. Available from: https://www.ncbi.nlm.nih.gov/books/NBK507256/

[20] Raikou, Vaia D.1; Kyriaki, Despina2,. The Relationship Between Concentrations of Magnesium and Oxidized Low-Density Lipoprotein and Beta2-microglobulin in the Serum of Patients on the End-stage of Renal Disease. Saudi Journal of Kidney Diseases and Transplantation 27(3):p 546–552, May–Jun 2016. | DOI: 10.4103/1319-2442.182396

[21] Rudloff S, Jahnen-Dechent W, Huynh-Do U. Tissue chaperoning-the expanded functions of fetuin-A beyond inhibition of systemic calcification. Pflugers Arch. 2022 Aug;474(8):949–962. doi: 10.1007/s00424-022-02688-6. Epub 2022 Apr 11. PMID: 35403906; PMCID: PMC8995415.

[22] Takata T, Isomoto H. The Versatile Role of Uromodulin in Renal Homeostasis and Its Relevance in Chronic Kidney Disease. Intern Med. 2024 Jan 1;63(1):17–23. doi: 10.2169/internalmedicine.1342-22. Epub 2023 Jan 15. PMID: 36642527; PMCID: PMC10824655.

[23] Kumar S, Sharma P, Arora K, Raje M, Guptasarma P. Calcium binding to beta-2-microglobulin at physiological pH drives the occurrence of conformational changes which cause the protein to precipitate into amorphous forms that subsequently transform into amyloid aggregates. PLoS One. 2014 Apr 22;9(4):e95725. doi: 10.1371/journal.pone.0095725. PMID: 24755626; PMCID: PMC3995793.

[24] Stiles LI, Ferrao K, Mehta KJ. Role of zinc in health and disease. Clin Exp Med. 2024 Feb 17;24(1):38. doi: 10.1007/s10238-024-01302-6. PMID: 38367035; PMCID: PMC10874324.

[25] Moskovskich A, Goldmann U, Kartnig F, Lindinger S, Konecka J, Fiume G, Girardi E, Superti-Furga G. The transporters SLC35A1 and SLC30A1 play opposite roles in cell survival upon VSV virus infection. Sci Rep. 2019 Jul 18;9(1):10471. doi: 10.1038/s41598-019-46952-9. PMID: 31320712; PMCID: PMC6639343.

[26] Qu YY, Guo RY, Luo ML, Zhou Q. Pan-Cancer Analysis of the Solute Carrier Family 39 Genes in Relation to Oncogenic, Immune Infiltrating, and Therapeutic Targets. Front Genet. 2021 Dec 2;12:757582. doi: 10.3389/fgene.2021.757582. PMID: 34925450; PMCID: PMC8675640.

[27] Yao JH, Ortega EF, Panda A. Impact of zinc on immunometabolism and its putative role on respiratory diseases. Immunometabolism (Cobham). 2025 Mar 5;7(1):e00057. doi: 10.1097/IN9.0000000000000057. PMID: 40051614; PMCID: PMC11882175.

[28] Jyothsna P, Suchitra MM, Kusuma Kumari M, Chandrasekhar C, Rukmangadha N, Alok S, Siddhartha Kumar B. Effect of Iron Deficiency Anemia on Glycated Albumin Levels: A Comparative Study in Nondiabetic Subjects with Iron Deficiency Anemia. J Lab Physicians. 2022 Oct 20;15(2):253–258. doi: 10.1055/s-0042-1757589. PMID: 37323601; PMCID: PMC10264113.

[29] Bilello JP, Cable EE, Isom HC. Expression of E-cadherin and other paracellular junction genes is decreased in iron-loaded hepatocytes. Am J Pathol. 2003 Apr;162(4):1323–38. doi: 10.1016/S0002-9440(10)63928-4. PMID: 12651624; PMCID: PMC1851226.

[30] Cleton MI, de Bruijn WC, van Blokland WT, Marx JJ, Roelofs JM, Rademakers LH. Iron content and acid phosphatase activity in hepatic parenchymal lysosomes of patients with hemochromatosis before and after phlebotomy treatment. Ultrastruct Pathol. 1988 Mar-Apr;12(2):161–74. doi: 10.3109/01913128809058215. PMID: 3363682.

[31] Jiang B, Liu G, Zheng J, Chen M, Maimaitiming Z, Chen M, Liu S, Jiang R, Fuqua BK, Dunaief JL, Vulpe CD, Anderson GJ, Wang H, Chen H. Hephaestin and ceruloplasmin facilitate iron metabolism in the mouse kidney. Sci Rep. 2016 Dec 19;6:39470. doi: 10.1038/srep39470. PMID: 27991585; PMCID: PMC5171654.

[32] Bergwik J, Kristiansson A, Allhorn M, Gram M, Åkerström B. Structure, Functions, and Physiological Roles of the Lipocalin α1-Microglobulin (A1M). Front Physiol. 2021 Mar 3;12:645650. doi: 10.3389/fphys.2021.645650. PMID: 33746781; PMCID: PMC7965949.

[33] Obeagu EI. Iron homeostasis and health: understanding its role beyond blood health - a narrative review. Ann Med Surg (Lond). 2025 May 21;87(6):3362–3371. doi: 10.1097/MS9.0000000000003100. PMID: 40486647; PMCID: PMC12140690.

[34] Tesfay L, Clausen KA, Kim JW, Hegde P, Wang X, Miller LD, Deng Z, Blanchette N, Arvedson T, Miranti CK, Babitt JL, Lin HY, Peehl DM, Torti FM, Torti SV. Hepcidin regulation in prostate and its disruption in prostate cancer. Cancer Res. 2015 Jun 1;75(11):2254–63. doi: 10.1158/0008-5472.CAN-14-2465. Epub 2015 Apr 9. PMID: 25858146; PMCID: PMC4454355.

[35] Kimita W, Ko J, Li X, Bharmal SH, Petrov MS. Associations Between Iron Homeostasis and Pancreatic Enzymes After an Attack of Pancreatitis. Pancreas. 2022 Nov-Dec 01;51(10):1277–1283. doi: 10.1097/MPA.0000000000002195. PMID: 37099767.

[36] Wang B, Zhang J, Song F, Tian M, Shi B, Jiang H, Xu W, Wang H, Zhou M, Pan X, Gu J, Yang S, Jiang L, Li Z. EGFR regulates iron homeostasis to promote cancer growth through redistribution of transferrin receptor 1. Cancer Lett. 2016 Oct 28;381(2):331–40. doi: 10.1016/j.canlet.2016.08.006. Epub 2016 Aug 11. PMID: 27523281.

